# Phasing of the ‘Wonderful’ Pomegranate Genome Using Haploid DNA Extracted from Pollen Grains

**DOI:** 10.64898/2026.07.13.738218

**Authors:** Giuseppe Lana, Ryan C. Traband, Sergio Pietro Ferrante, Mariano Resendiz, Lei Yu, Han Qu, Ethan Eurmsirilerd, Zhanao Deng, Mikeal L. Roose, Donald J. Merhaut, Taylor Beaulieu, Danelle K. Seymour, Frederick G. Gmitter, Zhenyu Jia, John M. Chater

## Abstract

The scientific and commercial interest in pomegranate (*Punica granatum* L.) cultivation has increased noticeably during the last two decades. Because of the high concentration of bioactive compounds and its promising nutraceutical properties, pomegranate has been defined as a functional food. In order to develop advanced genomic tools to improve pomegranate breeding program efficiency, we present the chromosome-scale and haplotype-resolved genome assembly of ‘Wonderful’, a pomegranate cultivar widely grown around the world. DNA isolated from diploid leaf tissues was sequenced using long read sequencing technology (PacBio and Nanopore), while DNA extracted from haploid pollen grains was sequenced using a short-reads platform (Illumina). Genomic data from 11 single haploid gamete cells were analyzed using the R package called *‘Hapi’* to phase the genome. The final genome assembly size was of 372.51 Mbp anchored to eight pairs of homologous chromosomes. The present study provides an insight on the adoption of an innovative and efficient approach for the assembly of haplotype-resolved genomes, which enables a higher resolution of DNA variant detection and offers the opportunity to investigate crossover events in single gamete cells during meiosis.

## Introduction

The consumption of pomegranate juice and arils has been linked to several potential human health benefits. Recent studies have highlighted the antioxidant and anti-inflammatory activities of this fruit, which have been suggested to prevent cardiovascular, neoplastic, neurological, metabolic, and intestinal disease [1]. The presence of phenolic acids, flavonoids, and other phenolic compounds like punicalagin are believed to possess anti-inflammatory and anti-aging properties. In addition, the natural compounds present in pomegranate may mitigate effects of chronic disease and metabolic disorders like diabetes, obesity, and insulin resistance. The interest of the medical community in this fruit has increased substantially in the last several years. Scientific reports have focused on clarifying the potential anticancer effects associated with the consumption of pomegranate products [2]. Increasing evidence indicates that polyphenols in pomegranate might positively modulate the function and composition of gut microbiota, thus playing a prebiotic role in the human intestine [3]. However, more clinical studies are needed to clearly understand pomegranate’s effects on human health.

Pomegranate production regions are faced with challenges such as long term-drought conditions and invasive pests and diseases [4, 5]. Increasing pomegranate biodiversity has been proposed as the primary strategy to reduce the risk of food system vulnerability related to monoculture and to increase the valorization of marginal land. Although ‘Wonderful’ represents the industry standard in the United States, several cultivars with desirable traits, such as low acidity and soft seededness, have been identified in the national pomegranate germplasm [6–8]. The high variability among pomegranate cultivars, combined with the crop’s capacity to adapt to a broad spectrum of soil and environmental conditions, represents a great resource for breeding and the development of new varieties that can expand cultivation to regions with more arid or more humid climates. Moreover, it will provide the opportunity to develop food products with improved sensory attributes and increased nutritional values [9, 10].

Many phasing technologies have been developed for analyzing diploid genomic data; however, reconstructing whole-chromosome haplotypes is still a challenging and expensive task. The several advantages provided by the knowledge of haplotypes, especially in the research fields of quantitative and evolutionary genetics, have stimulated the implementation of new approaches, such as constructing chromosome-scale haplotypes with gametes. Gamete cells represent an ideal material for inferring chromosomes because they are naturally haploid tissues and preserve the main parts of the allelic combinations. The usage of software for phasing chromosome-length haplotypes with gametes, like *Hapi* [11], helps to reduce the complexity of diploid-based phasing methods and minimize costs. In addition, it offers the chance to identify crossover events taking place during gamete formation, allowing the investigation of recombination frequency and the distribution of distances between crossover points [12] Figure 1.

**Figure 1.**
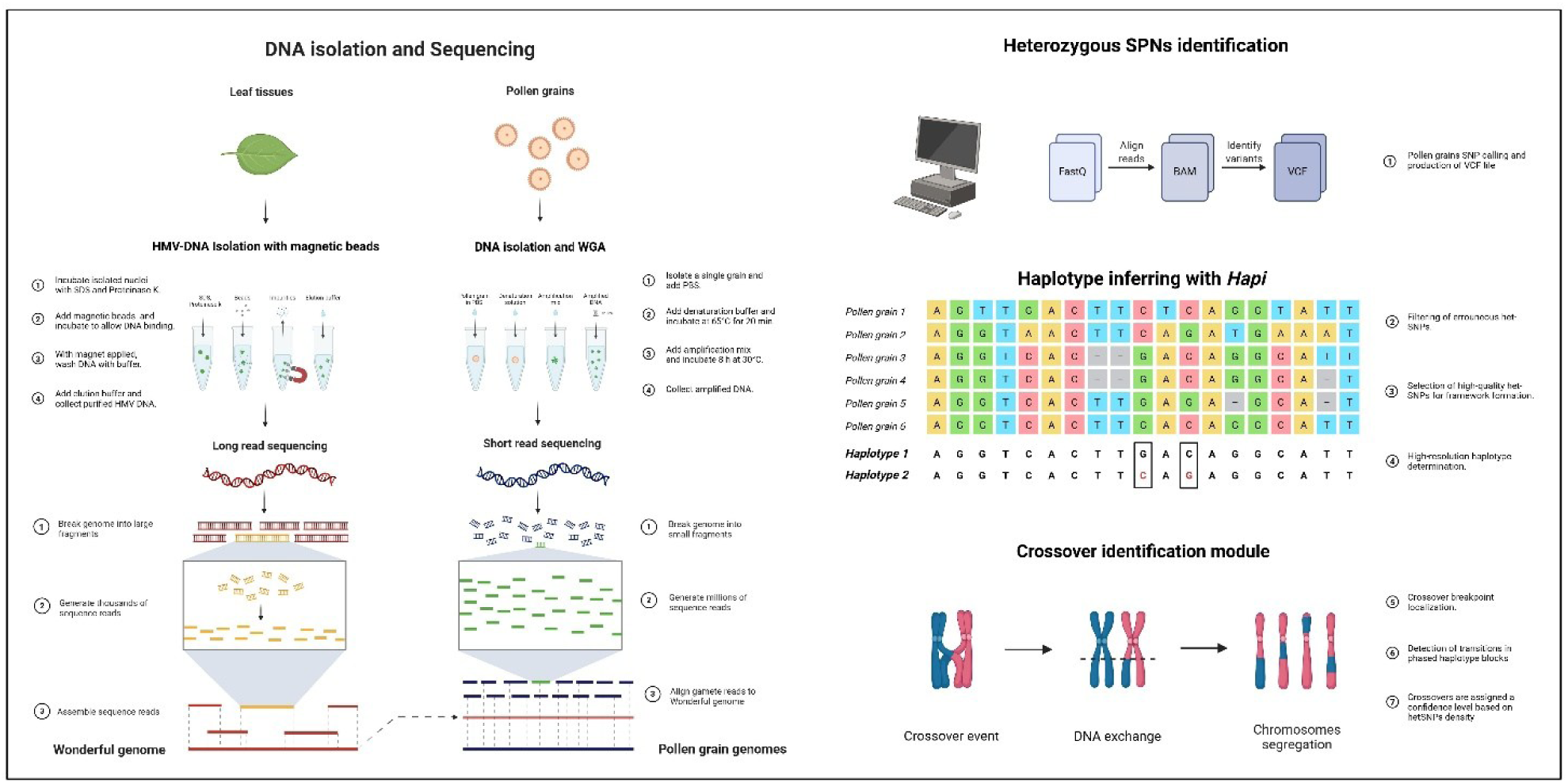
Workflow for Wonderful pomegranate genome phasing through pollen grain DNA analysis. This figure provides a simplified, conceptual illustration of the pipeline used to isolate genomic DNA from leaf tissue and single pollen grains, followed by high-throughput sequencing and bioinformatic analysis. DNA extraction protocols were optimized for both somatic and gametic tissues to ensure high-quality input for library preparation. Sequencing data were processed to identify heterozygous SNPs, which served as markers for haplotype phasing. Using the Hapi R package in RStudio, chromosome-length haplotypes were reconstructed from single-gamete genotypes, enabling precise inference of crossover events.

To provide a reliable genomic resource to support the identification of gene-trait associations, molecular marker development, variant discovery, and phylogenetics, we assembled and annotated *de novo* the ‘Wonderful’ pomegranate genome. The objective of this work was to make a high-resolution, chromosome-scale, and haplotype-resolved genome accessible for plant breeding using genomic pollen grain and Hi-C (high-throughput chromosome conformation capture) data. Transcriptome sequencing was included in the study for gene identification and annotation.

## Materials and Methods

### DNA isolation and sequencing

All the tissues utilized for the experiment were collected from the same *Punica granatum* L. tree cultivar ‘Wonderful’ located at the University of California Riverside Department of Agricultural Operations field facilities (GPS coordinates: 33.968703, -117.346451), which was originally propagated and sourced from the United States Department of Agriculture Agricultural Research Service National Clonal Germplasm Repository for Tree Fruit & Nut Crops & Grapes located in Winters, CA. After plants broke winter dormancy, young leaves were collected before sunrise. Tissues were immediately frozen in liquid nitrogen and stored at –80 °C until processed for nucleic acid isolation. High molecular weight DNA was extracted from the pomegranate leaves using an adapted version of the CTAB-based method, which combined two isolation protocols previously described by *Cheng et al*. and *Krizman et al.* [13, 14]. A detailed version of the protocol is available on Protocols.io [15]. The isolation was performed with little to no agitation of the DNA pellet to avoid any rupture of the DNA strands. Whole genome long read sequences were obtained by adopting PacBio HiFi CCS technology on a PacBio Revio platform provided with 25 million ZMW SMRT cell at fifty-fold coverage (Pacific Biosciences, CA, USA), which generated about 90 Gb of raw data. In addition, Oxford Nanopore sequencing (PromethION, Oxford Nanopore Technologies, UK) was performed to a depth of approximately one hundred-fold coverage to further enhance long-range contiguity and support assembly polishing. Hi-C proximity ligation data were generated using Proximo Genome Scaffolding Platform (Phase Genomics, WA, USA) to identify the physical collocation of the contigs into the chromosomes.

### RNA isolation and sequencing

Tissues utilized for the RNA isolation were collected from the same plant used for the DNA extraction. Six different types of tissues were selected to carry out the RNAseq: leaves, anthers, roots, bark, ovary and calix. RNA was isolated from tissue samples following a modified CTAB-based procedure previously developed by *Zarei et al*. [16]. Electrophoresis analysis was carried out to check RNA integrity on 1.0% agarose gel. Absorbance ratios 260/230 nm (≥ 1.90) and 280/260 nm (≥ 2.00) were acquired with Nanodrop One machine (Thermo Scientific). The RNA Integrity Number (RIN) for each sample, calculated by the company performing the RNA-seq, was equal to or above 7 for all libraries selected for sequencing. Polyadenylated RNA libraries were prepared and sequenced on the Illumina NovaSeq platform, generating approximately 150 million paired-end reads per sample following the manufacturer’s instructions.

### Pollen grain DNA isolation and sequencing

Genomic data from 11 single pollen grains were used for inferring chromosomal haplotype and obtaining a higher resolution on DNA variants detection. Anthers were collected from flowers just before anthesis. Pollen was harvested and dried at 30 °C and subsequently transferred to a 2 mL cryogenic vial and kept at –20 °C until processed. Single pollen grains were isolated under a stereomicroscope (10× - 40× zoom lens) using a sequencing pipette tip. Single pollen grains were transferred onto a custom micro-insert made from a 200 μl PCR tube, on a 1μl drop of BSA (0.5 mg/ml). Using a 40× zoom lens, the isolated pollen grains were checked to select only the one which had an intact structure.

Whole Genome Amplification (WGA) was carried out using a commercial kit (REPLI-g single cell kit, QIAGEN). Each micro-insert was placed inside a 200 ul PCR tube and pollen grains were crushed. 4 μl of PBS buffer and 3 μl of D2 buffer (provided with the kit) were added and pollen grains were incubated in a water bath at 65 °C for 20 minutes. The rest of the process followed the instructions provided by the manufacturer (QIAGEN protocol), except for the suggested WGA master mix. BSA was added to a final concentration of 1 mg/ml and incubated at 30 °C for 12 hours using a C1000 Touch Thermal Cycler (Bio-Rad), temperature of the lid was set to 70 °C. 16 SSR markers, two per each chromosome, were selected from a previously published study to evaluate the amplification performances of the DNA amplified from each pollen grain (Primer names: HvSSRT_81, HvSSRT_122, HvSSRT_222, HvSSRT_284, HvSSRT_375, HvSSRT_348, HvSSRT_497, HvSSRT_504, HvSSRT_592, HvSSRT_605, HvSSRT_713, HvSSRT_721, HvSSRT_773, HvSSRT_812, HvSSRT_826, HvSSRT_827) [17].

SSR amplification was performed on DNA extracted from single pollen grains using the same primer pairs previously validated in leaf tissue. PCRs were carried out in a total volume of 10 µL containing 1× PCR buffer, 2.0 mM MgCl₂, 200 µM of each dNTP, 0.2 µM of each primer, 0.5 U of Taq DNA polymerase, and 2 µL of whole-genome–amplified DNA. Reactions were run with an initial denaturation at 95 °C for 3 min, followed by 35 cycles of 95 °C for 30 s, locus-specific annealing temperature for 30 s, and 72 °C for 45 s, with a final extension at 72 °C for 5 min. PCR products were separated on 2% agarose gels and scored as successfully amplified when a clear, locus-specific band of the expected size was observed. For each pollen grain, we calculated the proportion of SSR loci that produced a clear, locus-specific amplicon. Because our panel included 16 SSR markers, grains were retained only if at least 12 of the 16 markers amplified successfully; grains below this threshold were discarded.

### Genome assembly and annotation

Two different sequencing techniques were adopted to achieve the best quality standards. Diploid DNA isolated from leaf tissues was sequenced using long reads sequencing (PacBio and Nanopore) and the short read (Illumina) technology was chosen to sequence haploid DNA amplified from pollen grains. The WGA DNA was used for Illumina sequencing to generate 150 bp single-end reads. The average minimum depth of coverage was about 5× for the 11 pollen grain samples. The ‘Wonderful’ *de novo* genome assembly was obtained by aligning both PacBio and Nanopore long reads with Hifiasm 0.25.0 software with default paramters [18]. The contigs obtained by the consensus sequence were assembled into chromosomes using Hi-C data. Raw Hi-C reads were mapped and filtered following the Phase Genomics mapping pipeline (https://phasegenomics.github.io/2019/09/19/hic-alignment-and-qc.html). Scaffolding of the draft assembly was conducted using YAHS v1.2a (Yet Another Hi-C Scaffolding tool) a command-line utility designed for constructing chromosome-scale assemblies by leveraging contact frequency data [19]. Hi-C contact map was generated using Juicer [20] and was visualized through Juicebox v1.11.08 [21]. The quality of the final assembly was assessed running BUSCO v5.7.0 (lineage embryophyta_odb10) and QUAST v5.2.0. To assess the continuity of the repetitive fraction, we calculated the LTR Assembly Index (LAI) using the LAI script included in LTR_retriever v2.9.0 [22, 23]. The analysis used the genome FASTA, the intact LTR-RT list generated during TE annotation, and the corresponding RepeatMasker output. Gene prediction and annotation was carried out following the Eukaryotic Genome Annotation Pipeline – External (EGAPx v1.0), a publicly accessible tool developed by NCBI (https://www.ncbi.nlm.nih.gov/refseq/annotation_euk/process/). The pipeline is designed to annotate genomes using a combination of protein homology, RNA-seq evidence, and ab initio prediction. It relies on Splign and Prosplign as alignment programs and Gnomon for gene prediction. Protein-coding genes were identified using *de novo* genome assembly and transcriptomic data (RNA-seq) as inputs. All software packages used in the assembly and evaluation workflow were executed with default parameters.

### Transposable elements identification

The Extensive de-novo Transposable Element Annotator (EDTA) v2.2.0 [24] was employed to identify and classify transposable elements (TEs) within the assembled ‘Wonderful’ genome using default parameters. The pipeline integrated structural and homology-based approaches to detect a broad spectrum of TE families, including long terminal repeat (LTR) retrotransposons, terminal inverted repeat (TIR) DNA transposons, and non-LTR elements. Divergence percentages for repeat landscape analysis were calculated using the Kimura 2-parameter (K2P) distance implemented in RepeatMasker, which estimates the sequence divergence between each TE copy and its consensus sequence. K2P values extracted from the RepeatMasker align output were used to generate the distribution of TE copies as a function of divergence (Figure 3), providing an approximation of relative insertion age.

**Figure 2.**
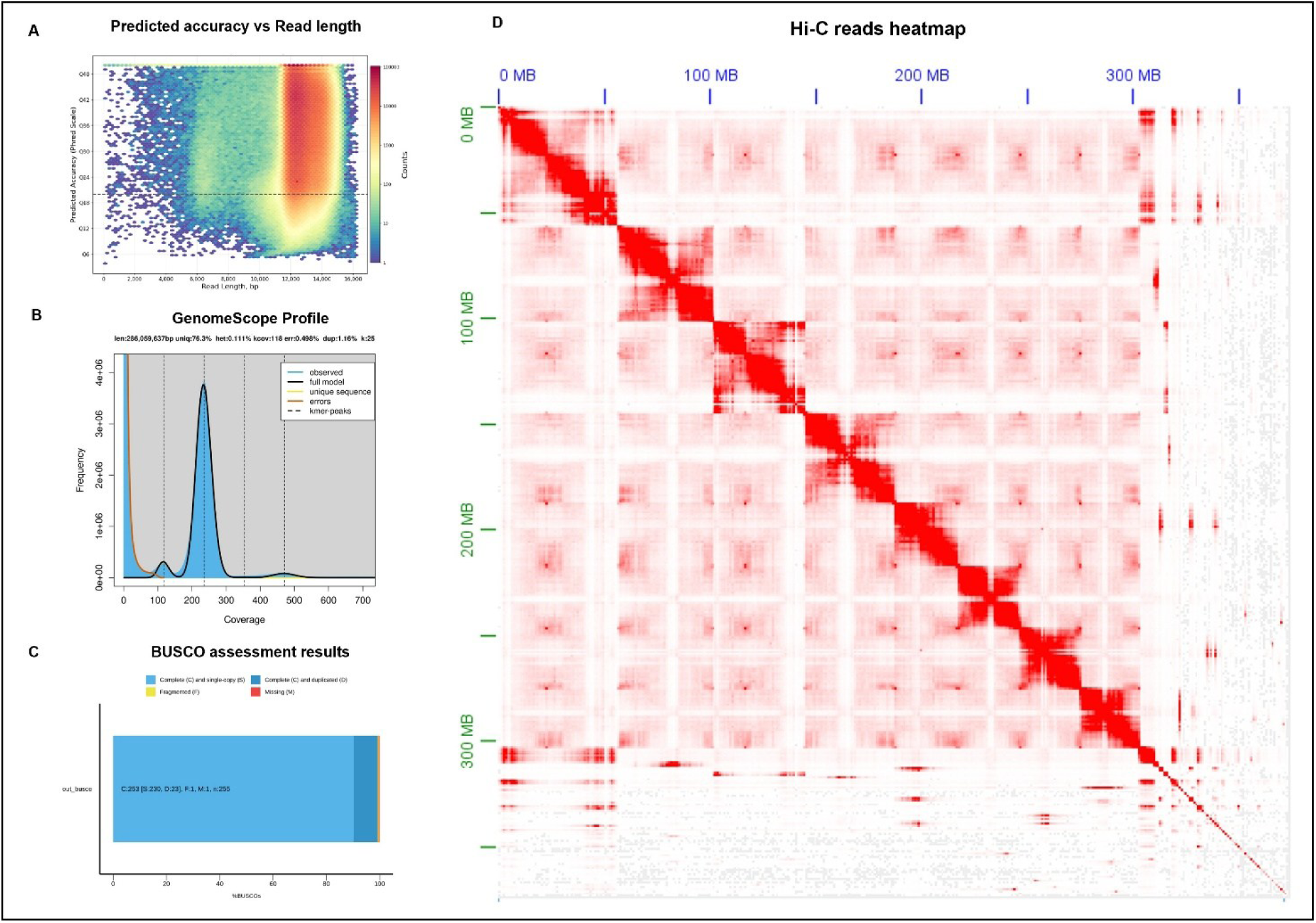
Quality assessment and chromosome-scale assembly validation. (A) Distribution of Phred quality scores across read lengths from raw sequencing data, indicating high base-calling accuracy throughout the read span. (B) The histogram displays the abundance of 25-mers across raw sequencing reads, with a prominent peak corresponding to the main coverage depth. (C) BUSCO analysis results based on the, showing the proportion of complete, duplicated, fragmented, and missing orthologs, reflecting the completeness and integrity of the assembled genome. (D) Hi-C contact map visualizing chromatin interactions across the genome, with strong diagonal signals confirming chromosome-scale contiguity and accurate scaffolding.

**Figure 3.**
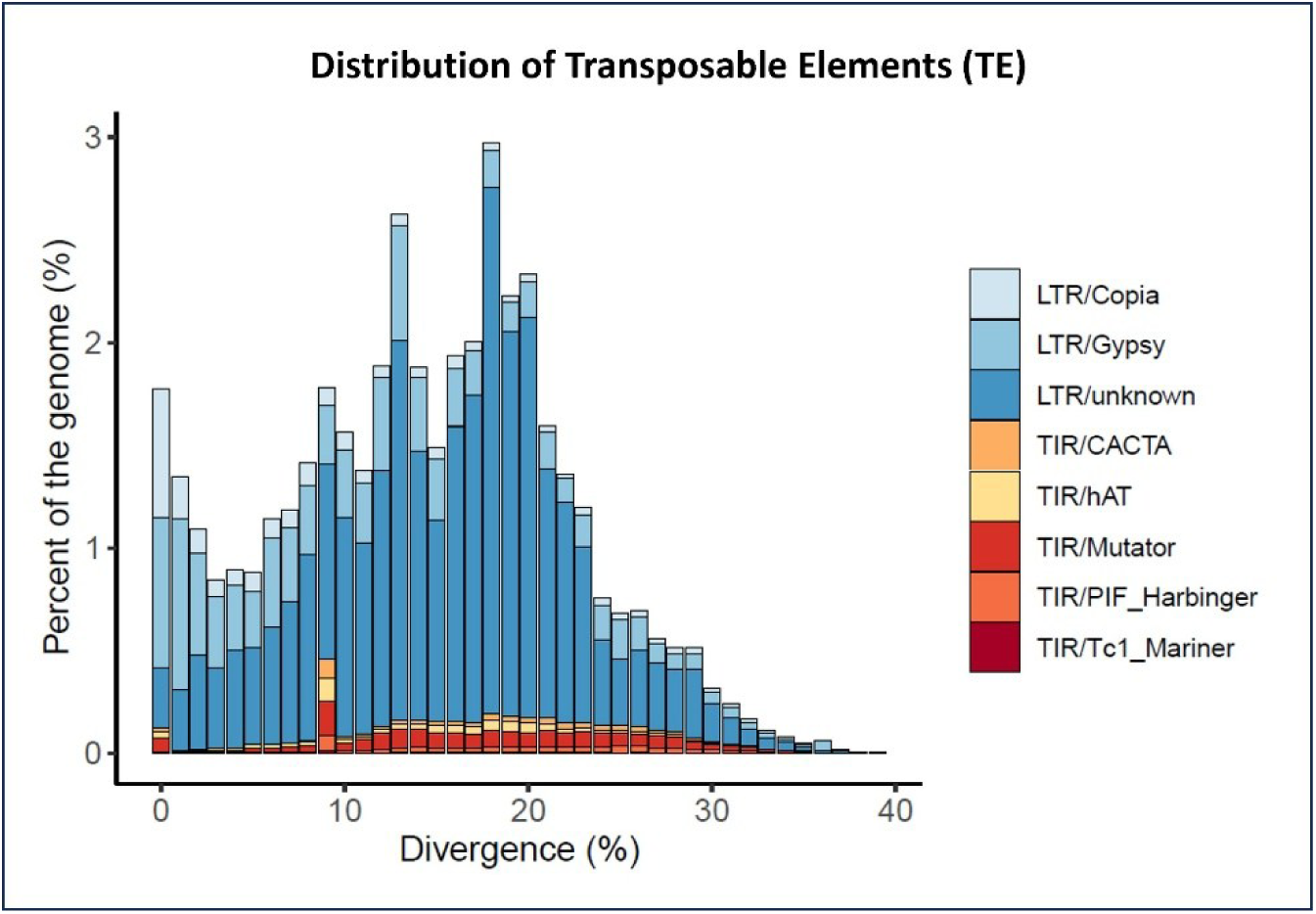
Histogram of transposable element abundance and genomic coverage in ‘Wonderful’ pomegranate. The graph illustrates the distribution of major TE superfamilies across the genome, highlighting the proportion of masked base pairs.

### Genome phasing

Raw Illumina reads obtained by sequencing DNA isolated from pollen grains were quality checked using FastQC v0.12.1. All the reads included in the analysis had a Phred score mean exceeding 30. Residual adaptor sequences and reads not compliant with quality standards were discarded using Trimmomatic v0.39 [25]. Haploid genomic data were mapped to the genome assembly using the Burrows-Wheeler Aligner v0.7.17 (BWA) [26]. Sam files created with BWA were converted into BAM files using SAMtools v1.18 and SNP calling was carried out. VCF files were obtained following the industry-standard GATK v4.4.0.0 Best Practices workflow for short-variant discovery (https://gatk.broadinstitute.org/hc/en-us). Variants were called for each pollen grain using HaplotypeCaller in gVCF mode, and the eleven resulting gVCFs were combined into a single joint-genotyped VCF. Because of the variable coverage typical of single-pollen genomes, hard filtering was applied to retain only high-confidence SNPs based on depth, mapping quality, genotype quality, and allele balance. The resulting filtered SNP set was then used to generate the input file for *Hapi* R Studio package [11]. The automatic phasing mode was utilized to run the package and identify the hetSNPs, while the crossover module allowed the identification of the crossover events.

### Synteny analysis

The pairwise whole-genome comparison between ‘Wonderful’ and ‘Tunisia’ sequences was conducted adopting a bioinformatics tool called SyRI v1.6.3 (Synteny and Rearrangement Identifier) [27]. The ‘Tunisia’ sequence was downloaded from GenBank where it was previously deposited under a Bioproject with accession number PRJNA565884 [28]. The two chromosome-level assemblies were previously aligned using Minimap2 v2.28 [29]. According to the suggestion of SyRI handbook, the alignment program was run on its preset for full genome/assembly alignment. The output file generated by SyRI was used to build the graphical representation of the genomic structure with plotsr.

## Results

### Genome assembly and quality assessment

The genome of ‘Wonderful’ pomegranate was sequenced with the single-molecule real-time technology (SMRT) at 50× coverage using high-fidelity (HiFi) long-read technology from Pacific Biosciences (PacBio). In addition, Oxford Nanopore long-read sequencing was performed on a high-output platform at approximately 100× coverage to further enhance long-range contiguity and support assembly polishing. The final genome assembly was 372.51 Mb, which corresponds to the 95.26% of the total estimated genome size. Assembly metrics, calculated using QUAST software [30], indicated that the assembly comprised 347 contigs, of which 318 were ≥ 25Mb and 253 were ≥ 50Mb, with the largest contig measuring ∼ 56Mb. The contig N50 and N90 were 42.34 Mb and 507,000 bp, respectively, while L50 and L90 were 4 and 33. The GC content was 42.26%, and there was a duplication ratio of 1.108 (Table 1).

**Table 1.**
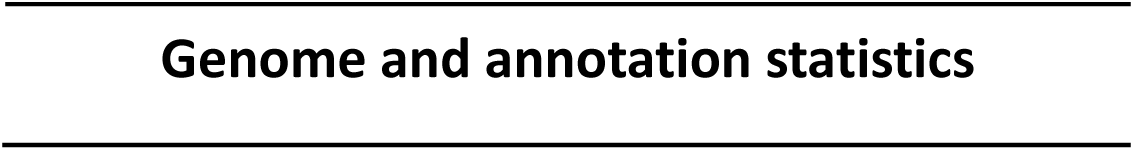
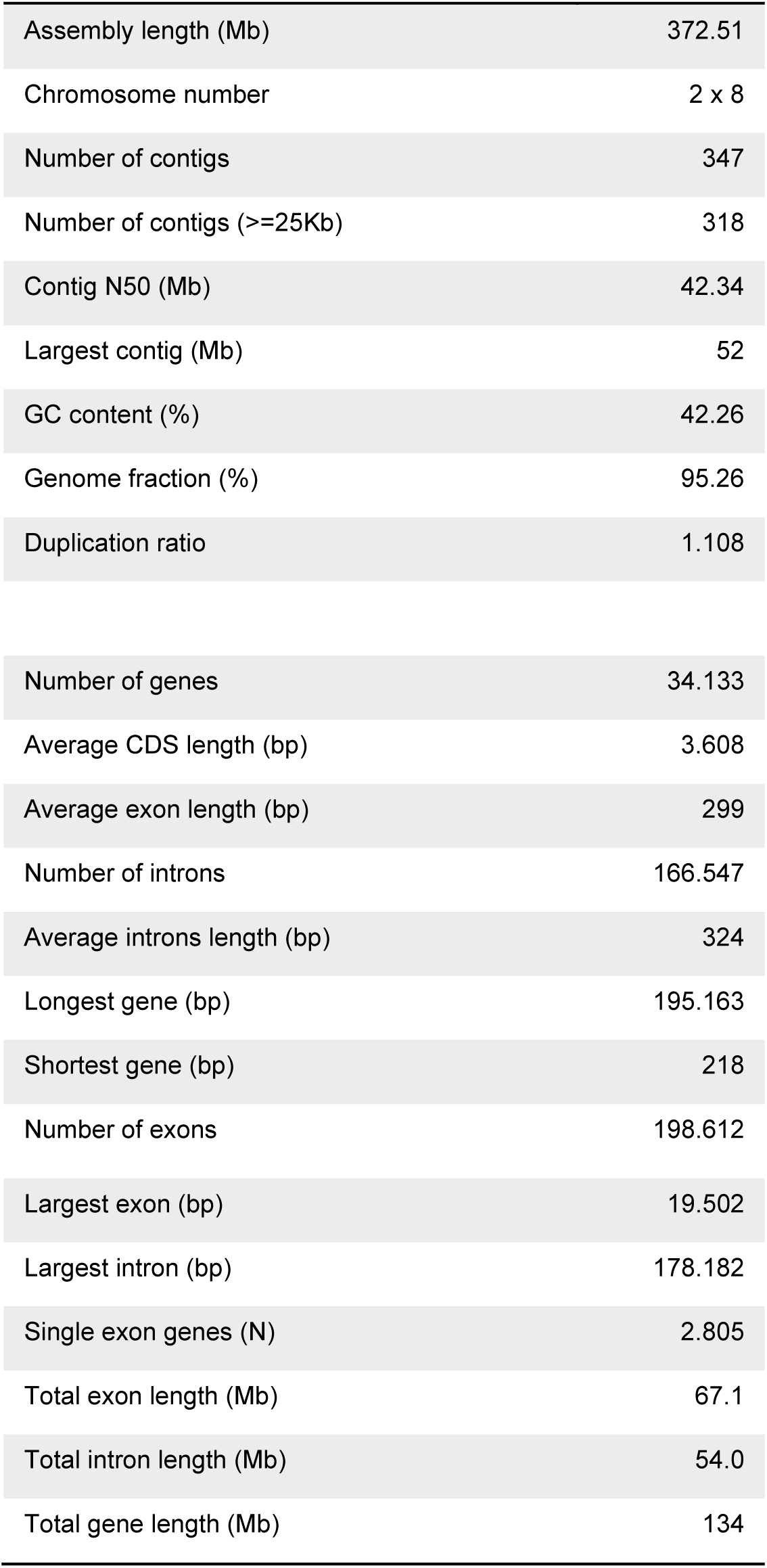
Summary of genome assembly and annotation statistics. This table presents key metrics from the genome assembly and structural annotation. Assembly statistics include total genome size, number of scaffolds, scaffold N50, and GC content. Annotation metrics report the number of predicted protein-coding genes, average gene length, exon count and some other key metrics.

|  |  |
| --- | --- |
| Assembly length (Mb) | 372.51 |
| Chromosome number | 2 x 8 |
| Number of contigs | 347 |
| Number of contigs ($\geq 25\text{Kb}$) | 318 |
| Contig N50 (Mb) | 42.34 |
| Largest contig (Mb) | 52 |
| GC content (%) | 42.26 |
| Genome fraction (%) | 95.26 |
| Duplication ratio | 1.108 |
| Number of genes | 34.133 |
| Average CDS length (bp) | 3.608 |
| Average exon length (bp) | 299 |
| Number of introns | 166.547 |
| Average introns length (bp) | 324 |
| Longest gene (bp) | 195.163 |
| Shortest gene (bp) | 218 |
| Number of exons | 198.612 |
| Largest exon (bp) | 19.502 |
| Largest intron (bp) | 178.182 |
| Single exon genes (N) | 2.805 |
| Total exon length (Mb) | 67.1 |
| Total intron length (Mb) | 54.0 |
| Total gene length (Mb) | 134 |

Proximity ligation data were employed to orient and order contigs into chromosome-scale scaffolds and assemble a reference-quality genome [31]. Hi-C data were generated using the Omni-C technology. After processing and normalizing the data, they were visualized and analyzed to observe the genome’s spatial organization. A chromatin interaction heatmap revealed that contigs were anchored to eight chromosomes (Figure 2, D).

To assess genome characteristics and sequencing quality, a *k*-mer frequency analysis was conducted using Jellyfish [32] and interpreted with GenomeScope [33]. Jellyfish was employed to count the frequency of 25-mers across the raw sequencing reads, generating a histogram that reflects the abundance of each *k*-mer (Figure 2, B). This distribution was then analyzed by GenomeScope, which modeled the data to estimate key genomic parameters including genome size, heterozygosity rate, repeat content, and sequencing error rate. The resulting profile revealed a 286 Mb genome with approximately 0.11% heterozygosity and 1.16% repetitive content. The distinct peak corresponding to the main *k*-mer coverage provided insight into the average sequencing depth and confirmed the overall structure of the dataset.

Assembly quality was evaluated using the Benchmarking Universal Single-Copy Orthologs (BUSCO) program [34], which revealed that 99.2% of the core eukaryotic genes were captured and complete. Notably, 90.2% of the genes were present as single copies, while 9.0% corresponding to 23 genes, were duplicated. Only 0.4% of the genes were fragmented, and just one gene was identified as missing (Figure 2, C). The assembly’s repetitive genome continuity was further supported by the LTR Assembly Index (LAI). Using the standard LTR_retriever workflow, we obtained a raw LAI of 21.12 and an adjusted LAI of 26.75. These values fall within the gold-standard range for LTR assembly continuity, highlighting the high structural integrity of the assembly across repeat-rich regions.

### Transposable elements annotation

The transposable element annotation revealed (Figure 3) a substantial proportion of repetitive content (51.50%), with LTR retrotransposons (35.28%), particularly Gypsy (8.70%) and Copia (2.40%) families, constituting the dominant TE class. The divergence landscape showed a prominent peak of LTR retrotransposons at low divergence (0–5%), consistent with recent transpositional activity, and additional peaks at higher divergence values (15–25%) indicative of older expansion waves. EDTA also generated a curated TE library and a comprehensive GFF3 annotation file, which can be used for downstream analyses such as genome masking, repeat landscape profiling, and comparative genomics. The default settings ensured a balanced trade-off between sensitivity and specificity, providing a reliable baseline for TE characterization in ‘Wonderful’. In addition to the overall TE composition, EDTA v2.2.0 provided detailed annotations of individual TE families, revealing lineage-specific patterns and structural diversity across our assembly. The annotation also captured fragmented and nested TE structures (24.18%), indicative of historical transpositional activity and genome remodeling. DNA transposons, including TIR elements such as Mutator (1.67%), hAT (0.75%), and CACTA (0.48%) families, were present at lower abundance. A remarkable abundance of non-TIR retrotransposons was detected, with 151,205 helitron elements comprising 9.89% of the genome. These results offer a foundational view of the repetitive landscape in ‘Wonderful’ pomegranate, providing insights into genome evolution, potential regulatory impacts, and targets for masking in downstream gene annotation workflows.

### Genome annotation

Transcript evidence was generated from several tissues to support genome annotation. mRNA-seq was carried out on six different plant tissues (leaves, anthers, roots, bark, ovary, and calix) and the sequences, after adequate preprocessing, were integrated into the gene prediction pipeline. A total of 34,133 mRNAs and 32,065 Coding DNA Sequences (CDS) were identified in our assembly, with an average coding sequence length of 3,608 bp and an average exon length of 299 bp. There were 166,547 introns present in the CDS with an average length of 324 bp. The longest gene detected measured 195,563 bp, while the shortest was only 218 bp long. The total number of exons detected in the CDS was 198,612, which translates to an average of 6.2 exons per gene. Additionally, 2,805 of the coding regions were identified as single exon genes. The total exon length was recorded to be 67.1 Mb, while the total introns length was 54.0 Mb, which corresponds to a total mRNA length of 134 Mb. The longest intron was 178,182 bp long, and the longest exon was 19,502 (Table 1).

### Inference of chromosomes

The genomic data obtained from the short reads sequencing (Illumina) of 11 independent pollen grains were used to adopt a gamete-based phasing method based on R studio package *Hapi* (Figure 1). The reads generated from the haploid tissues were aligned to the diploid genome assembly and variant calling was conducted to identify the heterozygous SNPs (hetSNPs). A total of 33,523 markers with potential genotyping errors were filtered out during data preprocessing to create a subset of high-quality hetSNPs. A total of 149,741 of hetSNPs were detected among the eight chromosomes and included in the chromosome frameworks that were used to construct the whole genome draft haplotypes. After each pollen grain was compared to the precursor framework to determine the gamete-specific variants, the final consensus high-resolution haplotype was created employing 138,890 hetSNPs (Figure 4). Out of the hetSNPs used for inferring the chromosomes, 56.9% had a high confidence level, about 38.5% were considered to have a fair level of confidence, and only 4.6% were rated as low confidence variants.

**Figure 4.**
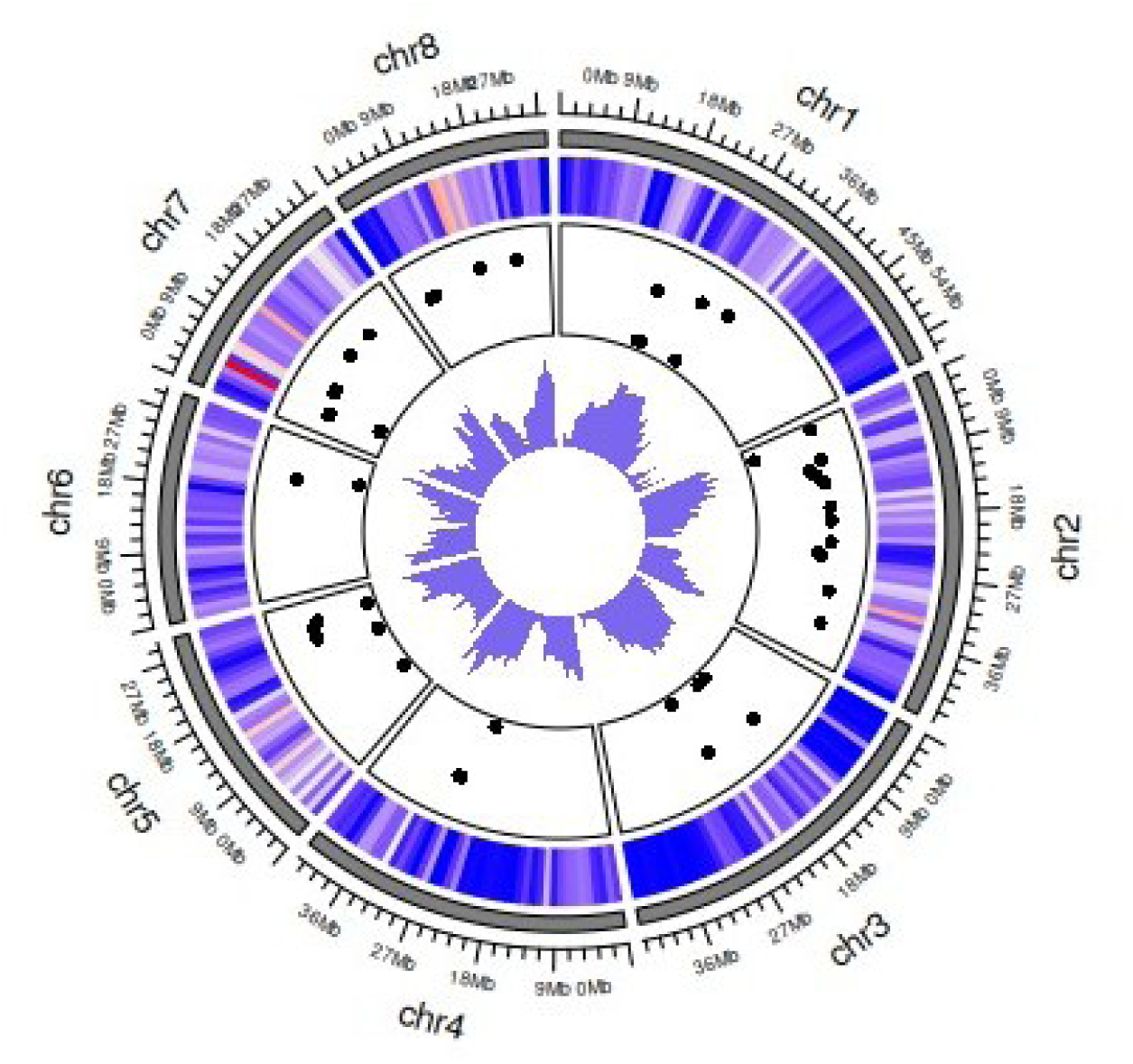
Circos plot summarizing genomic features across chromosomes. The outermost ring represents chromosome sizes, scaled proportionally. Inner tracks display (from outside to inside): heterozygous SNP density, crossover positions, and gene density. Colors in the SNP track represent SNP density from low (blue) to high (red). Crossover events are plotted as points along each chromosome; the x-axis corresponds to genomic coordinates, while the y-axis represents the distance between consecutive crossover events, providing a measure of crossover resolution. Gene density is shown as the number of annotated genes per fixed genomic window.

### Crossovers identification

The crossover analysis module included in Hapi R package provides a very important tool to investigate the recombination events that may occur in single gamete cells. The high-resolution chromosome-length haplotyping technique allowed for the identification and localization of 69 crossover events which took place among the 11 pollen grains utilized for the study. A frequency analysis of crossover events revealed notable variation in their distribution across chromosomal locations. Among the eight chromosomes examined, chromosome two exhibited the highest number of crossover events, with 15 occurrences, followed by chromosome 5 (13 events) and chromosome one (11 events). Chromosomes 3, 7, and 4 showed intermediate frequencies, with 7, 7, and 6 crossover events respectively, while chromosomes 6 and 8 were less frequently involved, with 3 and 4 events. These patterns may reflect underlying biological factors such as recombination hotspots, chromosomal architecture, or local sequence composition, and warrant further investigation in the context of meiotic dynamics and genome structure. The crossover resolution range spanned from 25 kb to 750 kb, with a higher density within the 250 kb interval (Figure 5). Additionally, the distribution of distances between adjacent crossovers was found to be within an interval < 20 Mb for over 75% of the identified crossovers. The graph of distances distribution between two adjacent recombination events showed a peak in correspondence of 3 Mb length.

**Figure 5.**
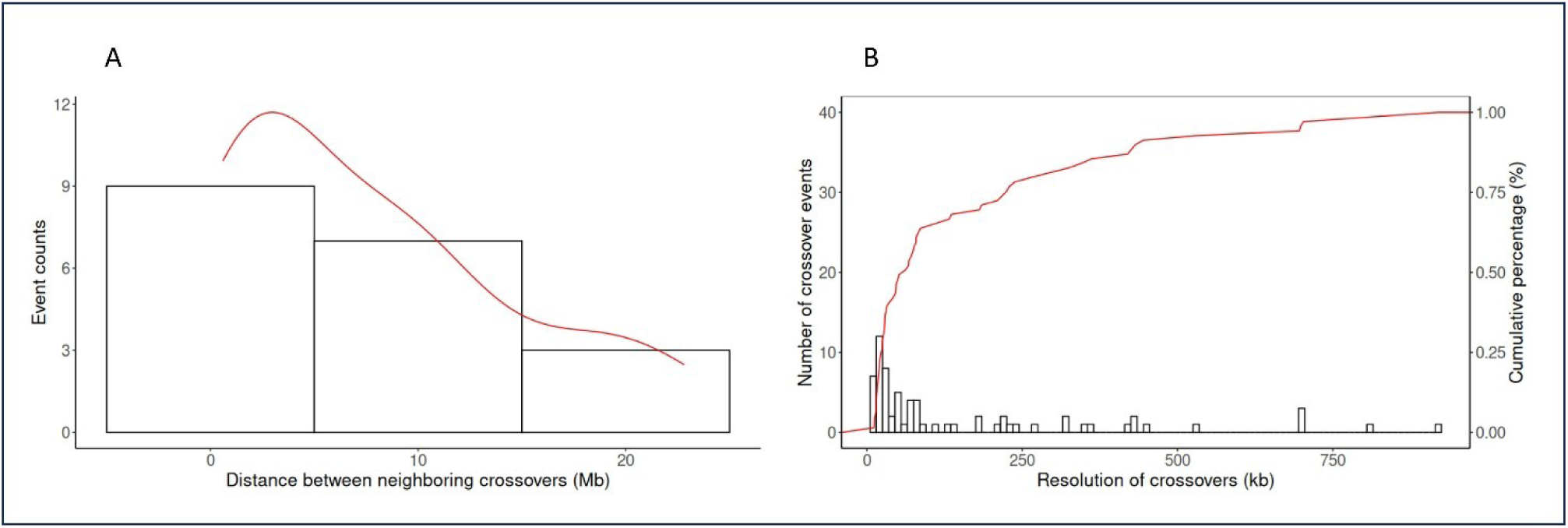
Analysis of inter-crossover distances and crossover resolution. (A) Histogram of inter-crossover distances within individual chromosomes, illustrating the spacing between consecutive crossover events. The red line represents a trend line summarizing the distribution. (B) Distribution of crossover resolution across the genome, defined as the physical span between flanking markers. In this panel, the red line represents the cumulative percentage and corresponds to the 1% value indicated on the right-hand scale.

### Synteny analysis

The ‘Wonderful’ and ‘Tunisia’ pomegranate genomes were compared to identify structural rearrangements and local sequence differences between the two assemblies. A syntenic analysis was conducted to detect inversion, translocations, or duplication events that might differentiate the two varieties. The results of the analysis revealed several rearrangements of the ‘Wonderful’ genome on chromosomes 1 and 3. According to the plot in Figure 6, several large blocks of the ‘Tunisia’ sequence had a different location in the ‘Wonderful’ assembly. Notably, a considerable portion of the sequence placed in the first half of chromosome 1 was translocated to the second part of the chromosome. Additionally, two inverted and two duplicated sections were included in the translocated regions.

**Figure 6.**
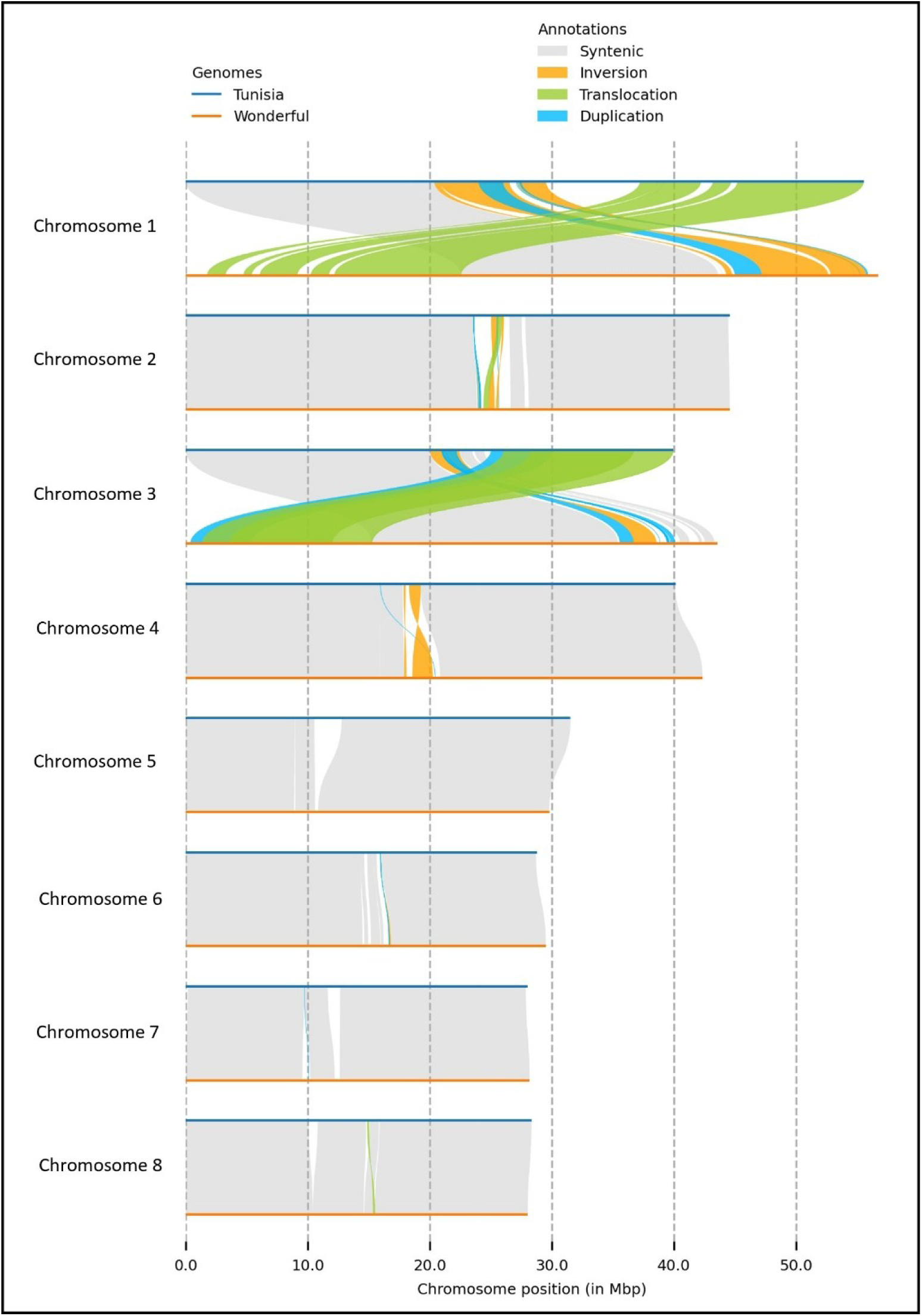
Synteny analysis between the ‘Tunisia’ and ‘Wonderful’ pomegranate genomes. Collinear blocks highlight conserved genomic regions between the two cultivars, revealing structural rearrangements, inversions, and potential translocation events. The alignment underscores both shared ancestry and cultivar-specific divergence, offering insights into genome evolution and breeding-relevant variation.

Although no remarkable structural variations were present on chromosomes 5, 6, 7 and 8, the comparison between the two genomes highlighted some non-syntenic blocks of variable size. Chromosome 5 was the only one where no structural variants were detected, while some very small rearrangements were found on chromosomes 6, 7, and 8. The sequences belonging to chromosomes 5, 6, 7, and 8 seemed to be the most conserved, since they were mostly syntenic. Similarly to chromosome 1, a large translocation was detected in chromosome 3. A large region located in the second half of the ‘Tunisia’ chromosome 3 was moved to the first part of the same chromosome in ‘Wonderful’. Two duplications and one inversion were included in these translocated regions. Some significant rearrangements were also highlighted in the central portions of chromosomes 2 and 4. Three small inverted regions, two duplications, and a translocated sequence were present in a non-syntenic region on chromosome 2. Two larger inversions and a smaller duplication event were observed in chromosome 4, in a region of lower synteny between these two genomes. Although these patterns could reflect structural variation based on the current data, and the available long-read and scaffolding evidence does not indicate misassembly, further analyses are required to determine whether these represent authentic translocations or assembly-related artifacts.

## Discussion

Although several efforts have been made to make available a high-quality whole genome assembly, none of the current published pomegranate genomes provides knowledge about haplotypes [28, 35–40]. A chromosome-scale genome assembly is a fundamental tool for both phylogenetic studies and crop improvement. The availability of genomic resources is indispensable for dissecting the genetic structure of the crop and understanding the relative evolutions of different species in comparison with other members of the family. Furthermore, it provides the means to identify genomic variants and associate them with various phenotypes. The previously published pomegranate genome assemblies are based on technology that supports individual SNP genotyping. While most genotyping studies are based on this methodology, modern genome assembly standards require access to chromosome-scale haplotype data. Haplotyping analysis offers increased resolution and accuracy of DNA sequences, enhancing the capacity for various genetic studies. Moreover, it can facilitate the identification of positive allelic combinations, especially in highly heterozygous or segregant populations [41].

Most available reference genome assemblies do not include separate sequences, preventing the distinction of haplotype regions inherited from maternal chromosomes from those inherited from paternal parent. This is because allele differences were ignored during the *de novo* assembly process, collapsing the genetic information of both parents into a unique consensus and chimeric sequence. Recent advancements in sequencing technology and haplotype phase generation promise more efficient identification of heterozygous loci [42, 43].

The determination for haplotyping phasing, which consists of identifying the alleles co-located on the same chromosome, can be based on experimental methods or computational approaches. The creation of double haploid genotypes and chromosome sorting are the most common laboratory-based methods. Sequencing double haploids provides the opportunity to assemble homozygous genomes, while separating chromosomes allows for the individual sequencing and assembly of the two haplotypes. Although these solutions are straightforward, they are very tedious and sometimes unfeasible due to technical limitations. For these reasons, computational phasing is the most frequently adopted technology to infer haplotypes [44].

Chromatin interaction detection with Hi-C represents the most accessible approach, successfully adopted for whole genome sequence scaffolding and haplotype phasing. However, this technology can generate contig orientation errors and haplotype switching because of the inability to accurately identify alleles that distinguish the homologous chromosomes inherited from the father and the mother [45, 46]. To address this challenge, a methodology called trio binning has been developed, where the whole-genome sequencing reads are separated into two haplotype-specific sets before the assembly. This approach relies on short reads generated from the parent lines to identify the genomic differences between the two haplotypes of the individual. Although efficient, this technique is not applicable when parents are unknown or not available, as in the case of ‘Wonderful’ [47].

The most widespread techniques currently used for inferring chromosome haplotypes are based on the analysis of diploid genetic materials, which may lead to partial and incorrect information due to the inability to perfectly distinguish and assemble maternal from paternal haplotypes [48]. For this reason, in the present work, we adopted an alternative phasing method based on the analysis of haploid tissues. Using gamete genetic material overcomes the technical limitations of diploid-based method by reducing the complexity of the inferring process [11, 12].

A recently published ‘Wonderful’ genome assembly by Slonimsky et al. provides an additional reference that complements existing genomic resources for *Punica granatum* (Slonimsky et al., 2025). Their population-level analyses revealed that domesticated pomegranates exhibit low heterozygosity and nucleotide diversity. The low heterozygosity reported for domesticated pomegranates, highlights an important limitation of diploid phasing approaches. When heterozygosity is sparse, Hi-C–based or read-based phasing becomes less informative, and distinguishing homologous chromosomes can be challenging. In contrast, the haploid pollen–based strategy used in the present study is not constrained by heterozygosity levels, because each pollen genome represents a single, fully phased haplotype. This makes pollen sequencing a robust and broadly applicable solution even in domesticated cultivars with reduced nucleotide diversity. Thus, the low-diversity patterns observed in the Slonimsky ‘Wonderful’ genome further underscore the utility of gamete-based phasing for resolving chromosome structure and allele-specific variation in pomegranate.

Meiotic recombination, specifically crossover formation between homologous chromosomes, is a fundamental process that ensures accurate chromosome segregation and promotes genetic diversity. During meiosis, crossovers facilitate the exchange of allelic variants, generating novel haplotypes that underpin phenotypic variation within populations [49]. For plant breeders, this mechanism is of strategic importance, as it enables the introgression of desirable traits and accelerates the development of improved cultivars. However, crossover events are not uniformly distributed across the genome; they tend to cluster in distal chromosomal regions while being suppressed in pericentromeric domains, thereby limiting the accessibility of certain loci to recombination-based breeding strategies [50]. Advances in high-resolution recombination mapping, haplotype phasing, and gamete genomics, particularly through the analysis of haploid pollen DNA, have enhanced the ability to detect and manipulate crossover landscapes [51]. These tools offer breeders greater precision in marker-assisted selection and facilitate the development of superior genotypes. Thus, meiotic crossovers represent not only a biological necessity but also a manipulable component in modern breeding programs [52, 53].

The genome assemblies of ‘Wonderful’, ‘Moshiliu’, and ‘Tunisia’ cultivars collectively represent a progressive refinement in pomegranate genomics, each contributing distinct strengths to breeding and functional studies [28, 54]. The ‘Wonderful’ genome, assembled using haploid pollen-derived data, achieved high phasing accuracy, a critical advancement for dissecting heterozygosity and structural variation in this crop. The phased assembly provided a valuable scaffold for allele-specific expression analysis and trait mapping in commercial breeding contexts.

In contrast, ‘Moshiliu’ stands out as the first T2T, gap-free assembly in pomegranate, spanning ∼366.71 Mb across eight fully resolved chromosomes, with complete centromere and telomere annotation. This structural clarity enabled the identification of key variants affecting anthocyanin biosynthesis, including a 37.2-kb translocation disrupting *PgANS* and a promoter repeat expansion in *PgANR*, both linked to peel pigmentation. The integration of multi-omics and GWAS across 146 accessions further positioned ‘Moshiliu’ as a reference for trait dissection and genome editing [54].

Meanwhile, the ‘Tunisia’ genome, assembled at ∼296.85 Mb, prioritized SSR marker development and polymorphism analysis, contributing over 365,000 perfect SSRs for population genetics and cultivar differentiation. Though less contiguous, ‘Tunisia’ offered insights into transposable element dynamics and soft-seed trait variation, particularly relevant for consumer-preferred phenotypes [28].

In addition to these resources, a recently published ‘Wonderful’ genome assembly (Slonimsky et al., 2025) provides a complementary reference based on PacBio long reads, Hi-C scaffolding, and genetic-map anchoring. These findings offer a domestication-focused perspective that complements the phased haplotype resolution achieved in the present study.

From a breeder’s perspective, ‘Wonderful’ provides a phased framework for allele-specific selection in elite cultivars; ‘Moshiliu’ delivers structural resolution for functional genomics and precision breeding; and ‘Tunisia’ offers a rich SSR toolkit for diversity analysis and traditional selection. Together, these assemblies form a complementary genomic triad that can advance *Punica granatum* L. improvement through distinct but synergistic lenses.

## Conclusion

In conclusion, the ‘Wonderful’ pomegranate genome assembly establishes a high-quality, phased reference that complements existing genomic resources such as ‘Moshiliu’ and ‘Tunisia’. By leveraging haploid pollen-derived data, the ‘Wonderful’ assembly enables accurate resolution of allele-specific variation, providing a robust foundation for comparative genomics and trait mapping. This resource facilitates the investigation of inter-cultivar genomic differences and supports the development of phenotype–genotype associations critical for breeding. The ‘Wonderful’ genome enhances the breeder’s toolkit for implementing advanced technologies such as molecular marker-assisted selection (MAS). This may reduce the time and resources required for the development of novel germplasm and accelerate the introgression of desirable traits, such as peel pigmentation, seed softness, and tree stress tolerance into commercially viable cultivars. The ‘Wonderful’ genome not only sets a new reference standard but also plays a strategic role in modernizing pomegranate breeding pipelines.

## Author Contributions

G.L. drafted the manuscript, performed data analysis and pipeline development, R.C.T., S.P.F., M.R., T.B., D.K.S. contributed to sample collection and DNA isolation, L.Y., H.Q., E.E. provided bioinformatic support and contributed to data interpretation, Z.D., M.L.R., D.J.M., F.G.G., Z.J., J.M.C. conceived and supervised the project. All authors reviewed and approved the submission of the final draft.

## Data Availability

The genome sequence data generated and analyzed during this study have been deposited in the National Center for Biotechnology Information (NCBI) under BioProject accession number **PRJNA1330292**. Individual sample metadata are available under BioSample accession numbers **SAMN51503288.** The assembled genome has been submitted to GenBank under accession number **JBRBXG000000000**. All datasets are publicly accessible and can be retrieved via the NCBI website (http://www.ncbi.nlm.nih.gov/bioproject/1330292). Metadata and associated annotations are provided in the supplementary materials.

## Declarations

### Funding declaration

This project was supported by the USDA Agricultural Marketing Service (AMS) MultiState Specialty Crop Block Grant through the California Department of Food and Agriculture project number 19-1043-002-SF.

### Ethics approval and consent to participate

Not applicable.

### Consent for publication

Not applicable.

### Competing interests

The authors declare no competing interests.

## Supplementary data

Additional supporting information can be found at the journal webpage on supplementary data section. **Table S1**. Hapi-derived heterozygous SNPs mapped across the Wonderful pomegranate genome for haplotype inference. **Table S2**: List of crossover events identified by Hapi on Wonderful pomegranate genome.

## Notes

### Competing Interest Statement

The authors have declared no competing interest.

## References

1. Caruso A, Barbarossa A, Tassone A, Ceramella J, Carocci A, Catalano A, et al. Pomegranate: Nutraceutical with promising benefits on human health. Applied Sciences (Switzerland). 2020;10:1–34. 10.3390/app10196915.

2. Pantiora PD, Balaouras AI, Mina IK, Freris CI, Pappas AC, Danezis GP, et al. The Therapeutic Alliance between Pomegranate and Health Emphasizing on Anticancer Properties. Antioxidants. 2023;12. 10.3390/antiox12010187.

3. Yin Y, Martínez R, Zhang W, Estévez M. Crosstalk between dietary pomegranate and gut microbiota: evidence of health benefits. Critical Reviews in Food Science and Nutrition. 2023. 10.1080/10408398.2023.2219763.

4. Luo Y, Hou L, Förster H, Pryor B, Adaskaveg JE. Identification of alternaria species causing heart rot of pomegranates in California. Plant Dis. 2017;101:421–7. 10.1094/PDIS-08-16-1176-RE.

5. Úrbez-Torres JR, Hand FP, Trouillas FP, Gubler WD. Pomegranate dieback caused by Lasiodiplodia gilanensis in California. Eur J Plant Pathol. 2017;148:223–8. 10.1007/s10658-016-1071-y.

6. Chater JM, Merhaut DJ, Jia Z, Mauk PA, Preece JE. Fruit quality traits of ten California-grown pomegranate cultivars harvested over three months. Sci Hortic. 2018;237:11–9. 10.1016/j.scienta.2018.03.048.

7. Chater JM, Merhaut DJ, Jia Z, Arpaia ML, Mauk PA, Preece JE. Effects of Site and Cultivar on Consumer Acceptance of Pomegranate. J Food Sci. 2018;83:1389–95. 10.1111/1750-3841.14101.

8. Chater JM, Santiago LS, Merhaut DJ, Jia Z, Mauk PA, Preece JE. Orchard establishment, precocity, and eco-physiological traits of several pomegranate cultivars. Sci Hortic. 2018;235:221–7. 10.1016/j.scienta.2018.02.032.

9. Adiba A, Razouk R, Charafi J, Haddioui A, Hamdani A. Assessment of water stress tolerance in eleven pomegranate cultivars based on agronomic traits. Agric Water Manag. 2021;243. 10.1016/j.agwat.2020.106419.

10. Díaz-Pérez JC, MacLean D, Goreta S, Workman S, Smith E, Sidhu HS, et al. Physical and chemical attributes of pomegranate (Punica granatum L.) cultivars grown in humid conditions in Georgia. HortScience. 2019;54:1108–14. 10.21273/HORTSCI13795-18.

11. Li R, Qu H, Chen J, Wang S, Chater JM, Zhang L, et al. Inference of chromosome-length haplotypes using genomic data of three or a few more single gametes. Mol Biol Evol. 2020;37:3684–98. 10.1093/molbev/msaa176.

12. Qu H, Li R, Yu L, Chen W, Feng Y, Jia Q, et al. IIIandMe: An Algorithm for Chromosome-scale Haplotype Determination Using Genome-wide Variants of Three Haploid Reproductive Cells. 10.1101/2022.12.07.519546.

13. Cheng Y-J, Guo W-W, Yi H-L, Pang X-M, Deng X. An Efficient Protocol for Genomic DNA Extraction From Citrus Species. 2003.

14. Križman M, Jakše J, Barièeviè D, Javornik B, Prošek M. Robust CTAB-activated charcoal protocol for plant DNA extraction

15. Resendiz M, Mar D, Mar P, Chammas A, Rose K, Tillman C, et al. high-molecular-weight-dna-extraction-of-pomegranate. ptotocols.io. 2025. 10.17504/protocols.io.j8nlkyobdg5r/v1.

16. Zarei A, Zamani Z, Mousavi A, Fatahi R, Karimi Alavijeh M, Dehsara B, et al. An Effective Protocol for Isolation of High-Quality RNA from Pomegranate Seeds. The Asian and Australasian Journal of Plant Science and Biotechnology . 2012;32–37 6 Special Issue 1.

17. Patil PG, Singh NV, Bohra A, Raghavendra KP, Mane R, Mundewadikar DM, et al. Comprehensive Characterization and Validation of Chromosome-Specific Highly Polymorphic SSR Markers From Pomegranate (Punica granatum L.) cv. Tunisia Genome. Front Plant Sci. 2021;12. 10.3389/fpls.2021.645055.

18. Cheng H, Concepcion GT, Feng X, Zhang H, Li H. Haplotype-resolved de novo assembly using phased assembly graphs with hifiasm. Nat Methods. 2021;18:170–5. 10.1038/s41592-020-01056-5.

19. Zhou C, McCarthy SA, Durbin R. YaHS: yet another Hi-C scaffolding tool. Bioinformatics. 2023;39. 10.1093/bioinformatics/btac808.

20. Durand NC, Shamim MS, Machol I, Rao SSP, Huntley MH, Lander ES, et al. Juicer Provides a One-Click System for Analyzing Loop-Resolution Hi-C Experiments. Cell Syst. 2016;3:95–8. 10.1016/j.cels.2016.07.002.

21. Durand NC, Robinson JT, Shamim MS, Machol I, Mesirov JP, Lander ES, et al. Juicebox Provides a Visualization System for Hi-C Contact Maps with Unlimited Zoom. Cell Syst. 2016;3:99–101. 10.1016/j.cels.2015.07.012.

22. Ou S, Chen J, Jiang N. Assessing genome assembly quality using the LTR Assembly Index (LAI). Nucleic Acids Res. 2018;46:e126. 10.1093/nar/gky730.

23. Ou S, Jiang N. LTR_retriever: A highly accurate and sensitive program for identification of long terminal repeat retrotransposons. Plant Physiol. 2018;176:1410–22. 10.1104/pp.17.01310.

24. Ou S, Su W, Liao Y, Chougule K, Agda JRA, Hellinga AJ, et al. Benchmarking transposable element annotation methods for creation of a streamlined, comprehensive pipeline. Genome Biol. 2019;20. 10.1186/s13059-019-1905-y.

25. Bolger AM, Lohse M, Usadel B. Trimmomatic: A flexible trimmer for Illumina sequence data. Bioinformatics. 2014;30:2114–20. 10.1093/bioinformatics/btu170.

26. Li H, Durbin R. Fast and accurate short read alignment with Burrows-Wheeler transform. Bioinformatics. 2009;25:1754–60. 10.1093/bioinformatics/btp324.

27. Goel M, Sun H, Jiao WB, Schneeberger K. SyRI: finding genomic rearrangements and local sequence differences from whole-genome assemblies. Genome Biol. 2019;20. 10.1186/s13059-019-1911-0.

28. Luo X, Li H, Wu Z, Yao W, Zhao P, Cao D, et al. The pomegranate (Punica granatum L.) draft genome dissects genetic divergence between soft- and hard-seeded cultivars. Plant Biotechnol J. 2020;18:955–68. 10.1111/pbi.13260.

29. Li H. New strategies to improve minimap2 alignment accuracy. Bioinformatics. 2021;37:4572–4. 10.1093/bioinformatics/btab705.

30. Gurevich A, Saveliev V, Vyahhi N, Tesler G. QUAST: Quality assessment tool for genome assemblies. Bioinformatics. 2013;29:1072–5. 10.1093/bioinformatics/btt086.

31. Belton JM, McCord RP, Gibcus JH, Naumova N, Zhan Y, Dekker J. Hi-C: A comprehensive technique to capture the conformation of genomes. Methods. 2012;58:268–76. 10.1016/j.ymeth.2012.05.001.

32. Marçais G, Kingsford C. A fast, lock-free approach for efficient parallel counting of occurrences of k-mers. Bioinformatics. 2011;27:764–70. 10.1093/bioinformatics/btr011.

33. Vurture GW, Sedlazeck FJ, Nattestad M, Underwood CJ, Fang H, Gurtowski J, et al. GenomeScope: Fast reference-free genome profiling from short reads. In: Bioinformatics. Oxford University Press; 2017. p. 2202–4. 10.1093/bioinformatics/btx153.

34. Manni M, Berkeley MR, Seppey M, Simão FA, Zdobnov EM. BUSCO Update: Novel and Streamlined Workflows along with Broader and Deeper Phylogenetic Coverage for Scoring of Eukaryotic, Prokaryotic, and Viral Genomes. Mol Biol Evol. 2021;38:4647–54. 10.1093/molbev/msab199.

35. Qin G, Xu C, Ming R, Tang H, Guyot R, Kramer EM, et al. The pomegranate (Punica granatum L.) genome and the genomics of punicalagin biosynthesis. Plant Journal. 2017;91:1108–28. 10.1111/tpj.13625.

36. Roopa Sowjanya P, Shilpa P, Patil GP, Babu DK, Sharma J, Sangnure VR, et al. Reference quality genome sequence of Indian pomegranate cv. ‘Bhagawa’ (Punica granatum L.). Front Plant Sci. 2022;13. 10.3389/fpls.2022.947164.

37. Yuan Z, Fang Y, Zhang T, Fei Z, Han F, Liu C, et al. The pomegranate (Punica granatum L.) genome provides insights into fruit quality and ovule developmental biology. Plant Biotechnol J. 2018;16:1363–74. 10.1111/pbi.12875.

38. Luo X, Shua Z, Zhao D, Liu B, Luo H, Chen Y, et al. Genome assembly of pomegranate highlights structural variations driving population differentiation and key loci underpinning cold adaption. Hortic Res. 2025;12. 10.1093/hr/uhaf022.

39. Patankar H V., Rivera LF, Alzahrani FO, Wing RA, Blilou I. A Chromosome level assembly of pomegranate (Punica granatum L.) variety grown in arid environment. Scientific Data . 2025;12. 10.1038/s41597-024-04337-2.

40. Slonimsky A, Lovky O, Harel-Beja R, Bar-Ya’akov I, Eshed R, Sherman A, et al. The pomegranate (Punica granatum L. cv. ‘Wonderful’) genome and P. protopunica shed light on pomegranate domestication. Is Daru a wild stock? BMC Genomics. 2025;26. 10.1186/s12864-025-11993-0.

41. Guk JY, Jang MJ, Choi JW, Lee YM, Kim S. De novo phasing resolves haplotype sequences in complex plant genomes. Plant Biotechnology Journal. 2022;20:1031–41. 10.1111/pbi.13815.

42. Zhang X, Wu R, Wang Y, Yu J, Tang H. Unzipping haplotypes in diploid and polyploid genomes. Computational and Structural Biotechnology Journal. 2020;18:66–72. 10.1016/j.csbj.2019.11.011.

43. Zhang W, Tariq A, Jia X, Yan J, Fernie AR, Usadel B, et al. Plant sperm cell sequencing for genome phasing and determination of meiotic crossover points. Nat Protoc. 2025;20:690–708. 10.1038/s41596-024-01063-2.

44. Browning SR, Browning BL. Haplotype phasing: Existing methods and new developments. Nature Reviews Genetics. 2011;12:703–14. 10.1038/nrg3054.

45. Xu Z, Dixon JR. Genome reconstruction and haplotype phasing using chromosome conformation capture methodologies. Brief Funct Genomics. 2020;19:139–50. 10.1093/bfgp/elz026.

46. Zeng Q, Zhou Z, He Q, Li L, Pu F, Yan M, et al. Chromosome-level haplotype-resolved genome assembly for Takifugu ocellatus using PacBio and Hi-C technologies. Sci Data. 2023;10. 10.1038/s41597-023-01937-2.

47. Delorean EE, Youngblood RC, Simpson SA, Schoonmaker AN, Scheffler BE, Rutter WB, et al. Representing true plant genomes: haplotype-resolved hybrid pepper genome with trio-binning. Front Plant Sci. 2023;14. 10.3389/fpls.2023.1184112.

48. Zhou Q, Tang D, Huang W, Yang Z, Zhang Y, Hamilton JP, et al. Haplotype-resolved genome analyses of a heterozygous diploid potato. Nat Genet. 2020;52:1018–23. 10.1038/s41588-020-0699-x.

49. Li X, Li L, Yan J. Dissecting meiotic recombination based on tetrad analysis by single-microspore sequencing in maize. Nat Commun. 2015;6. 10.1038/ncomms7648.

50. Luo C, Li X, Zhang Q, Yan J. Single gametophyte sequencing reveals that crossover events differ between sexes in maize. Nat Commun. 2019;10. 10.1038/s41467-019-08786-x.

51. Zhang W, Luo C, Scossa F, Zhang Q, Usadel B, Fernie AR, et al. A phased genome based on single sperm sequencing reveals crossover pattern and complex relatedness in tea plants. Plant Journal. 2021;105:197–208. 10.1111/tpj.15051.

52. Lian Q, Solier V, Walkemeier B, Durand S, Huettel B, Schneeberger K, et al. The megabase-scale crossover landscape is largely independent of sequence divergence. Nat Commun. 2022;13. 10.1038/s41467-022-31509-8.

53. Dreissig S, Fuchs J, Himmelbach A, Mascher M, Houben A. Sequencing of single pollen nuclei reveals meiotic recombination events at megabase resolution and circumvents segregation distortion caused by postmeiotic processes. Front Plant Sci. 2017;8. 10.3389/fpls.2017.01620.

54. Chen L, Wang H, Xu T, Liu R, Zhu J, Li H, et al. A telomere-to-telomere gap-free assembly integrating multi-omics uncovers the genetic mechanism of fruit quality and important agronomic trait associations in pomegranate. Plant Biotechnol J. 2025. 10.1111/pbi.70107.

