## Supplemental Table 2 for "Phasing of the ‘Wonderful’ Pomegranate Genome Using Haploid DNA Extracted from Pollen Grains"

***Table S2. List of crossover events identified by Hapi on the Wonderful pomegranate genome.*** *The table includes the gamete ID, chromosome number, start and end coordinates of each crossover event, the estimated crossover position, and its resolution.*

| Gamete | Chromosome | Start | End | Position | Resolution |
|---|---|---|---|---|---|
| W1 | 1 | 33282804 | 33302807 | 33292805 | 20002 |
| W1 | 2 | 18693342 | 18743319 | 18718330 | 49976 |
| W1 | 4 | 28078688 | 28109729 | 28094208 | 31040 |
| W1 | 5 | 18531594 | 18951343 | 18741468 | 419748 |
| W1 | 7 | 8260319 | 8275300 | 8267809 | 14980 |
| W1 | 7 | 8853327 | 8877704 | 8865515 | 24376 |
| W2 | 2 | 8916953 | 9101480 | 9009216 | 184526 |
| W2 | 2 | 11532721 | 11548417 | 11540569 | 15695 |
| W2 | 2 | 16218827 | 16243326 | 16231076 | 24498 |
| W2 | 6 | 23044613 | 23091159 | 23067886 | 46545 |
| W2 | 7 | 4371131 | 4422522 | 4396826 | 51390 |
| W2 | 8 | 14873219 | 15005588 | 14939403 | 132368 |
| W3 | 1 | 19217159 | 19235500 | 19226329 | 18340 |
| W3 | 1 | 29470775 | 29916167 | 29693471 | 445391 |
| W3 | 2 | 7168753 | 7179939 | 7174346 | 11185 |
| W3 | 2 | 25321249 | 25334937 | 25328093 | 13687 |
| W3 | 3 | 14432243 | 14468033 | 14450138 | 35789 |
| W3 | 3 | 16611500 | 16631600 | 16621550 | 20099 |
| W3 | 5 | 22777592 | 22800050 | 22788821 | 22457 |
| W3 | 5 | 23866395 | 24676317 | 24271356 | 809921 |
| W3 | 5 | 25391459 | 25620667 | 25506063 | 229207 |
| W3 | 6 | 21685418 | 21756813 | 21721115 | 71394 |
| W3 | 7 | 5693493 | 6396370 | 6044931 | 702876 |
| W3 | 7 | 15452835 | 15797952 | 15625393 | 345116 |
| W4 | 1 | 19391707 | 19412869 | 19402288 | 21161 |
| W4 | 2 | 25033826 | 25045590 | 25039708 | 11763 |
| W4 | 3 | 14459857 | 14533809 | 14496833 | 73951 |
| W4 | 6 | 23052540 | 23119532 | 23086036 | 66991 |
| W4 | 7 | 3519723 | 4216115 | 3867919 | 696391 |
| W5 | 3 | 13066524 | 13113422 | 13089973 | 46897 |
| W5 | 5 | 7657551 | 7736780 | 7697165 | 79228 |
| W5 | 5 | 18627394 | 18951343 | 18789368 | 323948 |
| W9 | 2 | 3048321 | 3065082 | 3056701 | 16760 |
| W9 | 3 | 24644554 | 24675098 | 24659826 | 30543 |
| W9 | 4 | 24315158 | 24376995 | 24346076 | 61836 |
| W9 | 5 | 18562677 | 18608467 | 18585572 | 45789 |
| W9 | 5 | 25235810 | 25250797 | 25243303 | 14986 |
| W9 | 8 | 4177095 | 4387298 | 4282196 | 210202 |
| W12 | 1 | 20150718 | 20375639 | 20263178 | 224920 |
| W12 | 1 | 29484312 | 29916167 | 29700239 | 431854 |
| W12 | 2 | 37557231 | 38088937 | 37823084 | 531705 |
| W12 | 3 | 24649983 | 24675098 | 24662540 | 25114 |
| W12 | 4 | 24349228 | 24376995 | 24363111 | 27766 |
| W12 | 5 | 22112867 | 22154740 | 22133803 | 41872 |
| W12 | 5 | 25235810 | 25250797 | 25243303 | 14986 |

| | | | | | |
|---|---|---|---|---|---|
| W13 | 1 | 27959121 | 28068665 | 28013893 | 109543 |
| W13 | 1 | 29484312 | 29916167 | 29700239 | 431854 |
| W13 | 2 | 8796809 | 8825753 | 8811281 | 28943 |
| W13 | 2 | 31581185 | 31667428 | 31624306 | 86242 |
| W13 | 4 | 24349228 | 24376995 | 24363111 | 27766 |
| W13 | 5 | 7700687 | 7729336 | 7715011 | 28648 |
| W22 | 1 | 19217159 | 19235500 | 19226329 | 18340 |
| W22 | 2 | 24870264 | 24889462 | 24879863 | 19197 |
| W22 | 5 | 18627394 | 18951343 | 18789368 | 323948 |
| W23 | 1 | 20150718 | 20375639 | 20263178 | 224920 |
| W23 | 2 | 9367279 | 10287215 | 9827247 | 919935 |
| W23 | 4 | 27423710 | 27502867 | 27463288 | 79156 |
| W23 | 5 | 7700687 | 7729336 | 7715011 | 28648 |
| W23 | 7 | 6158248 | 6396370 | 6277309 | 238121 |
| W23 | 7 | 20829904 | 20912118 | 20871011 | 82213 |
| W23 | 8 | 5285174 | 5421280 | 5353227 | 136105 |
| W24 | 1 | 20150718 | 20415816 | 20283267 | 265097 |
| W24 | 2 | 1193543 | 1374735 | 1284139 | 181191 |
| W24 | 2 | 3048321 | 3065082 | 3056701 | 16760 |
| W24 | 2 | 22679710 | 22691204 | 22685457 | 11493 |
| W24 | 3 | 16611500 | 16678710 | 16645105 | 67209 |
| W24 | 3 | 23509641 | 23871718 | 23690679 | 362076 |
| W24 | 4 | 24301223 | 24376995 | 24339109 | 75771 |
| W24 | 8 | 21356019 | 22054077 | 21705048 | 698057 |
