## Supplemental Table 1 for "Phasing of the ‘Wonderful’ Pomegranate Genome Using Haploid DNA Extracted from Pollen Grains"

***Table S1. Comprehensive list of heterozygous SNPs detected by Hapi in the Wonderful pomegranate genome assembly.*** *Each row represents a SNP with its chromosomal location (Chromosome, Position), reference and alternate alleles (Ref, Alt), and genotype calls across 13 pollen grains (W1–W13). Columns labeled W1–W13 correspond to individual haploid gametes used for haplotype phasing. Hap1 and Hap2 denote the inferred parental haplotypes. Total indicates the number of gametes with valid genotype calls at each SNP, Rate reflects the proportion of heterozygous calls among informative gametes, and Confidence (Conf) represents the phasing certainty assigned by Hapi.*

| Chr | Position | Ref | Alt | W1 | W12 | W13 | W2 | W22 | W23 | W24 | W3 | W4 | W5 | W9 | Hap1 | Hap2 | Total | Rate | Conf |
|---|---|---|---|---|---|---|---|---|---|---|---|---|---|---|---|---|---|---|---|
| 1 | 1812068 | G | A | NA | G | A | NA | G | G | A | NA | A | G | A | G | A | 8 | 1 | H |
| 1 | 1812655 | C | A | NA | C | A | NA | C | C | NA | NA | A | C | NA | C | A | 6 | 1 | H |
| 1 | 1813721 | T | C | NA | T | C | NA | NA | T | NA | NA | C | T | NA | T | C | 5 | 1 | H |
| 1 | 1819731 | G | A | NA | G | A | A | G | G | NA | NA | A | G | NA | G | A | 7 | 1 | H |
| 1 | 1819805 | G | C | NA | G | C | C | G | G | NA | NA | C | G | NA | G | C | 7 | 1 | H |
| 1 | 1824619 | G | A | NA | G | A | A | G | G | NA | NA | NA | G | NA | G | A | 6 | 1 | H |
| 1 | 1834547 | A | G | NA | A | G | NA | A | A | G | NA | G | A | NA | A | G | 7 | 1 | H |
| 1 | 1838200 | A | C | NA | A | C | NA | A | A | NA | NA | C | A | NA | A | C | 6 | 1 | H |
| 1 | 1838292 | C | T | NA | C | T | NA | NA | C | NA | NA | T | C | NA | C | T | 5 | 1 | H |
| 1 | 1839997 | C | T | NA | C | T | NA | C | NA | NA | NA | T | C | NA | C | T | 5 | 1 | H |
| 1 | 1850387 | A | G | NA | A | G | NA | A | NA | NA | NA | G | A | NA | A | G | 5 | 1 | H |
| 1 | 1853043 | C | A | NA | C | A | NA | C | NA | NA | NA | A | C | A | C | A | 6 | 1 | H |
| 1 | 1861493 | T | C | T | T | C | NA | T | NA | NA | NA | C | T | NA | T | C | 6 | 1 | H |
| 1 | 1862246 | T | G | T | NA | G | NA | T | T | NA | NA | G | T | G | T | G | 7 | 1 | H |
| 1 | 1863972 | A | G | A | A | G | NA | A | NA | NA | NA | G | A | G | A | G | 7 | 1 | H |
| 1 | 1866786 | T | A | T | T | A | NA | T | T | NA | NA | A | T | A | T | A | 8 | 1 | H |
| 1 | 1869992 | C | T | C | C | T | T | C | C | NA | NA | NA | C | T | C | T | 8 | 1 | H |
| 1 | 1871371 | G | A | G | G | A | NA | G | G | NA | NA | A | G | A | G | A | 8 | 1 | H |
| 1 | 1879113 | G | T | G | G | T | NA | NA | G | NA | NA | T | G | NA | G | T | 6 | 1 | H |
| 1 | 1879672 | A | G | A | A | G | NA | A | NA | NA | NA | G | A | NA | A | G | 6 | 1 | H |
| 1 | 1884466 | G | C | NA | G | C | NA | G | G | NA | NA | C | G | NA | G | C | 6 | 1 | H |
| 1 | 1886262 | C | G | NA | C | G | NA | C | C | NA | NA | G | C | NA | C | G | 6 | 1 | H |
| 1 | 1900810 | G | T | NA | G | T | NA | G | G | NA | NA | NA | G | T | G | T | 6 | 1 | H |
| 1 | 1903331 | C | A | NA | C | A | NA | C | C | A | NA | NA | C | NA | C | A | 6 | 1 | H |
| […] | | | | | | | | | | | | | | | | | | | |
| 8 | 27908711 | C | T | C | C | C | C | C | T | NA | C | NA | C | NA | T | C | 8 | 1 | F |
| 8 | 27912090 | T | C | T | T | NA | T | T | C | NA | T | NA | T | NA | C | T | 7 | 1 | F |
| 8 | 27912091 | G | A | G | G | NA | G | G | A | NA | G | NA | G | NA | A | G | 7 | 1 | F |
| 8 | 27922670 | G | A | G | G | NA | NA | G | A | NA | G | G | G | G | A | G | 8 | 1 | F |
| 8 | 27927099 | C | T | C | C | NA | C | C | T | NA | C | C | C | C | T | C | 9 | 1 | F |
| 8 | 27927374 | G | A | G | G | NA | G | G | A | NA | NA | G | G | G | A | G | 8 | 1 | F |
| 8 | 27930131 | A | G | A | A | NA | A | A | G | NA | A | NA | NA | A | G | A | 7 | 1 | F |
| 8 | 27934455 | G | A | G | G | G | NA | G | A | NA | G | G | NA | G | A | G | 8 | 1 | F |
| 8 | 27936959 | T | C | T | T | NA | NA | T | C | NA | NA | T | NA | T | C | T | 6 | 1 | F |
| 8 | 27948360 | T | C | T | T | NA | T | NA | C | C | T | NA | T | T | C | T | 8 | 1 | F |
| 8 | 27966495 | G | C | G | G | NA | G | NA | C | C | G | NA | G | G | C | G | 8 | 1 | F |
| 8 | 27972605 | T | C | T | T | NA | T | NA | C | C | T | NA | T | T | C | T | 8 | 1 | F |
| 8 | 27973477 | T | C | T | T | NA | T | NA | C | C | T | NA | T | T | C | T | 8 | 1 | F |
| 8 | 27986585 | G | A | G | G | G | G | G | A | A | G | NA | G | G | A | G | 10 | 1 | F |
| 8 | 27994554 | C | A | NA | C | NA | C | NA | A | A | C | C | C | NA | A | C | 7 | 1 | H |
| 8 | 27996758 | C | G | NA | NA | NA | NA | G | C | C | G | NA | G | G | C | G | 6 | 1 | H |
| 8 | 28006534 | C | T | NA | C | C | C | NA | T | T | C | C | NA | C | T | C | 8 | 1 | H |
| 8 | 28007526 | A | G | NA | A | A | A | NA | G | G | A | A | NA | A | G | A | 8 | 1 | H |
